# CNVmap: a method and software to detect and map copy number variants from segregation data

**DOI:** 10.1101/778753

**Authors:** Matthieu Falque, Kamel Jebreen, Etienne Paux, Carsten Knaak, Sofiane Mezmouk, Olivier C. Martin

**Affiliations:** GQE - Le Moulon, Université Paris-Saclay, INRA, CNRS, AgroParisTech, 91190 Gif-sur-Yvette, France; Department of Mathematics, An-Najah National University, Nablus, Palestine; GDEC, INRA, Université Clermont Auvergne, 63000 Clermont-Ferrand, France; KWS SAAT SE & Co.KGaA, Grimsehlstr. 31, 37574 Einbeck, Germany

**Keywords:** Copy number variation (CNV), Segregating populations, Allele frequency profiles, Non-Mendelian markers

## Abstract

Single nucleotide polymorphisms (SNPs) are widely used for detecting quantitative trait loci or for searching for causal variants of diseases. Nevertheless, *structural* variations such as copy-number variants (CNVs) represent a large part of natural genetic diversity and contribute significantly to trait variation. Over the past decade, numerous methods and softwares have been developed to detect CNVs. Such approaches are based on exploiting sequencing data or SNP arrays, but they bypass a wealth of information such as genotyping data from segregating populations, produced *e.g.* for QTL mapping. Here we propose an original method to both detect and *genetically map* CNVs using mapping panels. Specifically, we exploit the apparent heterozygous state of duplicated loci: peaks in appropriately defined genome-wide allelic profiles provide highly specific signatures that identify the nature and position of the CNVs. Our original method and software can detect and map automatically up to 33 different predefined types of CNVs based on segregation data only. We validate this approach on simulated and experimental bi-parental mapping panels in two maize and one wheat populations. Most of the events found correspond to having just one extra copy in one of the parental lines but the corresponding allelic value can be that of either parent. We also find cases with two or more additional copies, especially in wheat where these copies locate to homeologues. More generally, our computational tool can be used to give additional value, at no cost, to many datasets produced over the past decade from genetic mapping panels.

## Introduction

Single Nucleotide Polymorphisms (SNPs) are typically exploited *via* genotyping technologies, such as arrays or Genotyping by Sequencing, leading to high-density information on such polymorphisms. The wide availability of such tools explains why polymorphisms are principally characterized at this SNP level, even though it is known in many species that there is also a great deal of *structural* polymorphism across genomes. Generally one uses the terminology “structural variation” (SV) when there are inversions, translocations, insertions, deletions, or duplications involving segments of over 1000 base pairs. Identifying such SVs is a current challenge, rendered difficult by the so-called “complexity” of large genomes that involve many repetitive sequences, generally associated with transposable elements. Work on identifying SVs includes in particular searching for Copy Number Variations (CNVs) (Redon *et al*. 2006) and Presence/Absence Variations (PAVs) (Alkan *et al*. 2011). In CNVs, a gene or segment of DNA is present in different numbers of copies in two genomes. CNVs have been discovered in numerous species, and in particular in mammals (Guryev *et al*. 2008; Conrad *et al*. 2010). In PAVs, a gene or DNA segment is present in one genome and missing in the other. It has been claimed that more nucleotide bases are affected by SVs than by SNPs (Zhang *et al*. 2009). Both CNVs and PAVs are associated with phenotypic variations and diseases (Beckmann *et al*. 2007; Zhang *et al*. 2009), making them a focus of much current research. A large number of approaches have been used to detect structural variations. Perhaps the oldest is based on competitive hybridizations to oligonucleotide probes as occurs in comparative genomic hybridization (CGH) arrays (Beló *et al*. 2009; Springer *et al*. 2009). A somewhat different technique is used in SNP arrays where individual samples are genotyped (no competition between samples). In such arrays, a fluorescence intensity is associated with each allele of a SNP; by following these intensities for successive SNPs along the genome, it is possible to identify regions in which the signal is anomalously low for all alleles, indicative of a PAV, or in which some intensities are a few fold higher than expected, indicative of a CNV (Colella *et al*. 2007; Cooper *et al*. 2008). Later approaches involve exploiting sequencing read depth (Bailey *et al*. 2002; Yoon *et al*. 2009; Alkan *et al*. 2009), unexpected mappings of read pairs (Chen *et al*. 2009; Pang *et al*. 2010), split reads (Mills *et al*. 2006; Ye *et al*. 2009), and sequence-based reassembly (Zerbino and Birney 2008; Chaisson *et al*. 2009; Simpson *et al*. 2009; Li *et al*. 2010). Each approach has its advantages (Alkan *et al*. 2009; Sudmant *et al*. 2010) and continues to undergo optimization (Zare *et al*. 2017). Our work takes a novel approach, different from all that we listed above, and gives “a second life” to SNP genotyping technologies in the specific context of populations in segregation. Indeed, SNP arrays have been available at low cost for several years now and so have been used rather extensively to genotype segregating populations. Such populations are typically constructed for mapping quantitative trait loci (QTLs) for fundamental research or for breeding programs. Interestingly, such technologies provide, in the context of segregating populations, information on structural variation between the founding parents used to build the population. However, that information is hidden within the markers showing non-Mendelian segregation patterns, markers that generally are discarded early-on in the linkage mapping analyses. The present work provides a methodology for inferring certain types of SVs based on those non-Mendelian markers in bi-parental mapping populations. We will stress the CNV cases, because in such situations the previously unknown copies of the region can also be *mapped* based on the linkage disequilibrium analyzed from the segregation data. Indeed, our approach exploits particular *profiles* of allele frequencies arising along the genome, somewhat analogously to what is done in genome-wide association studies. However, in our case, instead of working with the genotype at a single marker at a time, we work with compound genotypes involving multiple markers. The profiles built in this way exhibit peaks at the loci where the extra copies arise and provide signatures allowing the identification of the type of SV involved. These additional loci should not be too close in genetic distance to the reference locus because, as we shall see, it is the recombinations between these loci in the population that lead to the characteristic signal.

Any marker that deviates strongly from Mendelian behavior in a segregating population points to a potential SV. The simplest context in which to understand how to exploit a non-Mendelian signal is to consider the case of doubled-haploid (DH) plant populations in segregation. Indeed, in that type of population where homozygous individuals are derived from single meiotic products by chromosome doubling, all of the Mendelian markers should show full homozygosity while SVs and more specifically CNVs will lead some individuals to appear as being heterozygous at markers involved in rearrangements. Therefore, we begin our study here using a doubled haploid (DH) population consisting of 625 maize individuals; we refer to it as the GABI population. We will also show how our approach can be used in the context of RIL (Recombinant Inbred Line) and IRIL (Intermated Recombinant Inbred Line; (Lee *et al*. 2002)) populations. The maize IRIL population studied in this paper is the IBM (Ganal *et al*. 2011) mapping population. We also study a RIL wheat population (Choulet *et al*. 2014; Rimbert *et al*. 2018), hereafter referred to as WHEAT. The markers in these RIL and IRIL populations have residual heterozygosity because the number of generations of selfing used is not so large. Such blocks of markers with residual heterozygosity might be expected to swamp any SV signal; interestingly this turns out not to be true. We will (1) demonstrate the efficiency of our approach on our three populations that include two situations with this potential problem, (2) show that the allele frequency profiles are fully compatible with the inferred CNVs by comparing those profiles to the ones produced from simulated populations, and (3) compute *p*-values associated with the H_0_ hypothesis that the observed profiles occurred by chance in the absence of SV. As a first illustration, we apply our method and software to the GABI maize DH population, revealing striking signals of duplications and triplications, the corresponding copies arising on a *common* chromosome not more often than by chance. Then we examine the case of recombinant inbred lines (RILs or IRILs); as expected, having residual heterozygosity makes the problem more challenging but our method generalizes well. Indeed, in the IBM and WHEAT experimental populations, we are able to identify unambiguous signatures of copy number variations. Interestingly, in the WHEAT data set we find a large number of triplicated loci that involve the homeologous chromosomes. Finally, we assess the candidate CNVs in the IBM population using reference genome sequences of the parents.

## Materials and Methods

### Aim and design of the study

The locus-specificity of genetic markers used for genotyping, be they PCR-based or array-based, mostly relies on oligonucleotides hybridizing to sequences flanking the SNP of interest. In the case of duplicated regions, such oligonucleotides may find alternative targets, messing up the interpretation of the observed raw data (mostly fluorescence intensities at two wavelengths), which usually assumes a single target locus. This results in apparent non-Mendelian behaviours of some markers, which are usually filtered out from data sets before using them for mapping or QTL analyses. The goal of this work was to exploit these types of data sets to infer markers involved in SVs and genetically map the previously unknown copies, providing a software package to do so automatically. This software is provided as a R package named CNVmap, provided in Supplementary File S1 (see Supplementary File S6 for installation). More detailed explanations about the functionalities and procedures of this software are provided in the package *via* the embedded documentation of the associated functions.

In our approach, we focus on non-Mendelian markers to provide and test appropriate hypotheses, interpreting the observed segregations as CNV events polymorphic between the parents used to generate segregating populations. The main signature of such events being apparent heterozygous calls, we therefore worked with segregating populations in which the individuals are almost fully homozygous, like doubled-haploid or recombinant inbred populations. In such populations, high levels of heterozygous calls in some markers strongly point to possible CNVs containing those markers. Because some systematic or random errors in the genotyping process can also lead to unexpected heterozygotes, we extended our approach to a joint analysis of each candidate marker with its local allelic context, which provides multiple and unambiguous signatures that make up strong evidence for the reality of the event. Moreover, this approach (implemented in our software) also provides the genomic localization of the other loci involved in the CNV.

### Populations and segregation data used

The population mainly used in this study is a Doubled Haploid (DH) maize population called GABI (Presterl *et al*. 2007), from which genotyping data were kindly provided by KWS SAAT SE & Co.KGaA (Einbeck, Germany). The GABI population contains 625 DH lines, which were genotyped using the Illumina MaizeSNP50 array (Ganal *et al*. 2011). Second, we also studied the maize IBM population (Ganal *et al*. 2011) which is is an Intermated Recombinant Inbred Line (IRIL) population obtained by intermating F2 individuals for four generations to accumulate recombination, before beginning the selfing generations used to fix the material (Lee *et al*. 2002). The IBM population contains 239 RILs, which were genotyped using the same SNP array and cluster file as for GABI. Lastly, we studied a wheat population consisting of, 406 F6 individuals derived from a cross between Chinese Spring and Renan; these were genotyped using the TaBW280K SNP array (Rimbert *et al*. 2018). Marker segregation data for populations GABI, IBM, and WHEAT are respectively provided in Supplementary Files S2, S3, S4.

### Linkage map construction

Our method and software to detect CNVs requires prior knowledge of the order and genetic position of the markers. The genetic maps used to analyze the GABI and IBM segregation data were thus first obtained using a seriation approach implemented in R scripts calling functions from the CartaGene software (de Givry *et al*. 2004), as described in Ganal et al. (Ganal *et al*. 2011). The genetic map used to analyze the segregation data of the WHEAT population were produced using the software MSTmap (Wu *et al*. 2008) with the following default parameters: population type: RIL6; distance function: Kosambi; cut-off: 0.00000000001; map dist.: 15; map size: 2; missing threshold: 0.20; estimation before clustering: yes; detect bad data: yes; objective function: ML (Rimbert *et al*. 2018). Linkage map data for populations GABI, IBM, and WHEAT are respectively provided in Supplementary Files S2, S3, and S4.

### Raw data filtering

In DH populations, each normal marker should be homozygous in every offspring. The possible calls for any marker are then “A” (the allele attributed to parent 1) or “B” (the allele attributed to parent 2). It is important to keep in mind that a call of a SNP is the result of an elaborate identification process which is not 100% reliable so that one cannot exclude a low proportion of errors, leading for instance to some small proportion of “H” calls (heterozygous, which should almost never happen in a DH population if there is no genotyping error) or “-” for missing data if the calling is too ambiguous. In practice, we do see heterozygous markers in spite of each individual being homozygous. Such cases might be erroneous calls or indicative of regions belonging to SVs. Beyond possible errors in the calls for certain markers, some individuals of the population may themselves be corrupt (e.g. through pollen contamination). A strong indication of this is if an offspring has an anomalously large number of markers that are called “H”. We thus apply some quality filtering on the data sets, both at the level of the individuals (e.g. we cast out individuals having too high heterozygozity rates) and at the level of markers. In the R code, the user can change the thresholds for such filterings, for instance based on minor allele frequency or genotype frequencies (see all parameter descriptions in the R package).

### Defining the Mendelian and candidate markers

Given the markers passing the previous test, we now divide them into three classes: “Mendelian”, “candidate”, and “other”. Our procedure for defining the first or second class of markers is based on forcing them to be respectively “typical” and “atypical” for some statistic while the “other” markers are all the ones that do not pass these tests. Our first statistic is the fraction of individuals that are heterozygous. We require that this fraction be in the bottom X_H_ percentile for “Mendelian” and in the top Y_H_ for “candidate” markers. We do the same thing (but with different thresholds) using the fraction of individuals that are called missing for that marker. Our use of percentiles has the advantage that it automatically takes into account the properties of the population, such as the low numbers of heterozygotes in DH populations and the significantly larger numbers arising in RILs that have not reached fixation. Clearly, as the threshold Y_H_ is lowered, the number of candidate markers will increase so if one wants to find as many events as possible pointing to SVs it is good to not take Y_H_ too large. In contrast, the potential number of Mendelian markers is quite substantial so it is not a problem to be rather stringent for the value of X_H_.

### Automatic detection of peaks in the allele profiles

For each candidate marker M* we identify its flanking Mendelian markers ML and MR from which we identify the individuals in the population belonging to each of six associated 3-marker genotype classes (see Supplementary File S5). Then for each of these six classes we compute the corresponding genome-wide allele profile using the *Mendelian* markers only, each marker leading to a frequency defined as the number of individuals carrying the A allele divided by the number of individuals carrying either the A or B alleles. These six genome-wide allele profiles along chromosomes are then analyzed for occurrences of peaks. Roughly a peak can be defined *via* a region on the genome in which the allele frequency curve has a pointed shape and approaches very close to 0 or 1. In practice, to avoid being sensitive to noisy or erroneous data, we get rid of outliers by a first filter. That means producing a first smoothed version of the allele curve using splines (*smooth.spline()* function in R) and throwing out the data points that are outliers with respect to that curve. Second, a new smoothed curve is generated using the remaining markers. Then all regions for which this new smoothed curve is close enough to 0 or 1 are identified. A linear fit of the data (outliers excluded) is performed on each side of the putative peak to determine its expected position, and also to assess the quality and slope of the linear regression on both sides of the peak (or on one side only if the peak is at the extremity of a chromosome, or close to another peak). Then the list of all peaks for all six classes are compared to see whether peaks co-localize. This leads to a list of peaks (genetic positions on the genome) with each peak being called as “present” or “absent” for each of the six allele frequency curves. Note that if a class contains no individuals it is just ignored (see Supplementary File S5). Furthermore, when there are few individuals in a class, the associated allele frequency curves are noisy and thus will have peaks by chance. We thus only consider classes having a minimum number of individuals, this minimum being determined so that by the Bonferroni test one has a false discovery rate for peak detection that is 5% under the hypothesis that there is no structural variation present.

### Automatic assignment to a type of CNV event

Once all peaks have been detected, for the associated locus the presence or absence of peaks or troughs for the list of 6 different 3-marker genotype classes of individuals was encoded with a 6-character string. The list of these strings (one per locus) provided the observed signature of the event. This signature was then compared by the software with a list of 33 predefined signatures (details provided as Supplementary File S5), and in case of a match between the observed signature and a predefined one, then the event was assigned to the corresponding type. The predefined signatures were based on theoretically expected patterns arising from CNVs involving additional copies at one or two loci. Such signatures depend on the allelic content at these different loci, leading us to introduce below a schematic notation for CNV events.

### Nomenclature used for the different types of CNVs

In the following, a CNV involving in parent 1 X doses of the genomic region of interest and Y doses in parent 2 will be referred to as a “X:Y CNV” (X and Y being equal to 1, 2, or 3). Moreover, each CNV category is encoded as a string of 2 to 3 groups of 3 characters, there being one group per locus, each separated by an underscore. Each group contains the parental alleles separated by a slash, so the result takes the following form: A group of 3 characters specifies the alleles carried at the considered locus by parents 1 and 2, in that order, separated by a slash, A being the reference allele of M* in parent 1 and B being the reference allele of M* in parent 2. The different groups are further concatenated using the underscore as a separator for the successive loci: locus1(P1/P2)_locus2(P1/P2)_locus3(P1/P2). The first group is always encoded “A/B” and indicates the reference locus, located at the position where the candidate marker was initially mapped. Further groups indicate the different additional loci carrying copies of the region targeted by the candidate marker. So for instance for a 2:1 CNV of type A/B_B/-the Parent 1 has two copies of the considered genomic region but the copy at the second locus carries the allele B, while the Parent 2 has only one copy.

### Analyzing candidates based on missing data

Our method is based on the detection of candidate markers for which the number of H calls is anomalously high, followed by an analysis of each associated genome-wide allelic profile. However, when for completeness we tried to analyze allelic profiles for *all* markers, we discovered clear CNV-like signatures for some markers with little or no H calls but with large numbers of missing data. In such cases, instead of the AHA or BHB 3-marker genotypic class, the peaks were observed on the allelic profiles associated with A-A or B-B classes suggesting that one had a CNV but where, for unknown reasons, the H calls for the candidate marker were transformed into missing data calls. So we specified in the software the signatures that would arise from having H calls be erroneously modified and denoted them by adding a suffix to their putative CNV type. The suffix was “|Hm” when (part of) the expected H calls have been turned into missing data calls (with probability *p_Hm_*), and “|HmHa” respectively “|HmHb” when the expected H calls have been turned partly into missing data calls (with probability *p_Hm_*) and partly into A respectively B homozygote calls (with probabilities *p_Ha_* respectively *p_Hb_*).

To simulate such events, we first estimated the probabilities *p_Hm_*, *p_Ha_*, and *p_Hb_* from the data for the marker considered based on the allelic profiles at the inferred peaks. We then simulated the CNV event as explained below. Finally, we introduced systematic “errors” to the resulting candidate marker genotype depending on the type of the putative CNV event with its suffix. Specifically, we randomly transformed the H calls into missing data, A, or B calls according to the probabilities *p_Hm_*, *p_Ha_* and *p_Hb_*.

### Producing simulated datasets

We produced simulated data staying as close as possible to the experimental population parameters, keeping the same marker positions on the genetic maps of each chromosome and the same population size. We simulated the exact same scheme of crossings as the one used to produce the experimental populations, implementing *in silico* crossovers that can arise in each marker interval during each meiosis based on the experimental genetic map. Crossover interference was also implemented using the Gamma model (McPeek and Speed 1995) whose parameter *nu* can be set as a parameter in our software (for typical values of *nu* in maize, see (Falque *et al*. 2009)). This implementation of interference proved to be important for having comparable peak width between experimental and simulated profiles. To simulate any particular CNV hypothesis, we implemented into the parental genomes the associated duplications or triplications of the marker M* of interest, using positions of loci inferred from the analysis of the actual experimental population. The corresponding modified parental genomes thus had extra fictitious markers each tagged with the parental allelic value (and thus independent of the CNV hypothesis). For instance in the case of a duplication in Parent A but with opposite allele (CNV of the type A/B_B/-), the extra marker had nevertheless allele A in parent A and allele B in parent B. Then the scheme of crossings was simulated based on these modified parental genomes (note that the genetic map was also modified but just by the inclusion of the extra markers at their inferred positions). Lastly, the individuals in the resulting population were “genotyped” *in silico*. For the markers that were not involved in the CNV, this was straightforward. However, to genotype an individual for the marker M*, it was necessary to take into account allelic values not only at M* but also at the extra copy or copies of the marker, to mimic the fact that oligonucleotides used for genotyping M* would hybridize on all copies. This was where the actual CNV hypothesis intervened because the “raw” genotypes at each extra marker as produced by the simulation had to be reinterpreted using the CNV allelic content. Specifically, the call of the marker M* had to be changed to H if and only if the reference locus was not already H and both A and B alleles were present in the reinterpreted individual when considering M* and all of its copies. As an illustration of this rule, consider again the CNV of the type A/B_B/-. The only situation requiring that a call of M* be changed to H is when the raw genotype is A at the first locus and also A at the second. In practice we apply such “transformation” rules using successively each of the extra copies of the marker, each time testing whether the genotyping should be changed to H. Once that is done, the extra copies are removed from the data set and only the original markers and associated modified calls are used as input to the analysis program, leading to production of corresponding genome-wide allelic profiles based on these simulated data sets. A good agreement between profiles produced from the experimental and simulated data sets then provides strong support for the hypothesized CNV.

### Calculation of p-values associated with the hypothesis H_0_ of no structural variation

Although having the simulated profiles allows one to get a feeling for whether a proposed CNV is plausible through consistency between theory and experiment, it is appropriate to also compare to the null hypothesis H_0_ whereby there is no CNV and the marker M* is present in only one copy in both parents. Under that hypothesis, the additional peaks in the experimental allelic profiles are simply due to stochasticity in the segregation, a situation that will be a problem whenever relatively few individuals contribute to these profiles. The CNVmap package provides a test of H_0_ in the form of a *p*-value that is computed as follows. Let M* be the considered marker that is a candidate for belonging to a region involved in a CNV. In our first step we identify within the whole population two sub-classes of individuals: the ones for which the flanking (Mendelian) markers of M* are both called as A alleles, and the ones for which those markers are both called as B alleles. Not all individuals fall within one of these classes, so for instance if for an individual one flanking marker is heterozygous, or if one is A and the other is B, then the individual is not further considered. Within each sub-class, the errors (heterozygous and/or missing data calls) under H_0_ are random. Thus the second step of our procedure is to produce a simulated dataset by shuffling the calls of M* separately in each of the two sub-classes of individuals defined in the first step. Under H_1_ (presence of a CNV) the M* calls are correlated with the calls at the second locus, while under H_0_ (M* is single-locus in both parents) there is no such second locus. The third step is to apply our analysis pipeline to this shuffled dataset and identify the peaks in the allelic profiles. The second and third step are repeated a large number of times (this number is specified by the user and computed *via* parallelization). Lastly, the *p*-value for rejecting the hypothesis H_0_ is obtained from the fraction of the shufflings that lead to having additional peaks in the allelic profiles.

### Use of parental genome sequences for validating CNVs predicted from the IBM population

To provide independent validation of CNVs detected with our software, we examined the whole-genome sequence assemblies of the two parents (B73 and Mo17) used to produced the IBM population. First, for each non-Mendelian marker M* identified with our software as being located in a 1:2 or 2:1 CNV, we extracted from the B73 sequence *three* regions 201 bp long (100bp before and 100bp after the SNP) flanking not only the marker M* (indicating the reference locus) but also each of the two markers (M_left_peak_ and M_right_peak_) delimiting the second locus (identified automatically in our software *via* the corresponding fitted peak positions). To do that, we used the V2 version of the B73 genome assembly (AGPv2 RefGen_v2 https://www.maizegdb.org/genome/genome_assembly/B73%20RefGen_v2) because the physical coordinates of the MaizeSNP50 SNPs are given on that V2 version (Ganal *et al*. 2011). Then, for our CNV validation, we BLASTed these three 201bp sequences against the B73 AGPv4 RefGen_v4 Maize genome assembly (the most recent available assembly https://www.maizegdb.org/genome/genome_assembly/Zm-B73-REFERENCE-GRAMENE-4.0) using default parameters. We then considered that the presence of a second copy in B73 was validated if one of the high-scoring pairs (HSP) returned by BLAST for the M* flanking region was on the same chromosome as the second locus and was included in the confidence interval of that locus (based on coordinates of HSPs obtained when BLASTing M_left_peak_ and M_right_peak_ flanking regions). Presence of the second locus in B73 is expected in 2:1 CNVs but not in 1:2 CNVs. We proceeded similarly for testing the presence of both loci in the Mo17 parent, except that we first extracted the three 201bp regions of Mo17 corresponding to M*, M_left_peak_ and M_right_peak_ markers by BLASTing the 201bp B73 regions against the Mo17 genome (https://www.maizegdb.org/genome/genome_assembly/Zm-Mo17-REFERENCE-CAU-1.0). We then BLASTed those 201bp Mo17 sequences against the Mo17 genome assembly. The second locus is then expected to be present in Mo17 in 1:2 CNVs but not in 2:1 CNVs.

### Availability of data and material

All data generated or analysed during this study are included in this published article and its supplementary information files (software provided as Supplementary File S1 and data sets provided as archives in Supplementary Files S2, S3, and S4). All Supplementary Figures, Tables, and Files have been uploaded to FigShare.

## Results

### Genome-wide allele frequency profiles identify the loci involved in CNVs

#### Strikingly clean signatures for 1:2 or 2:1 CNVs

What should be expected in a segregating population if only one of the parents has a marker duplicated? The simplest situation is schematically represented in Fig. 1A where in parent 1 (with alleles denoted “A”) a DNA segment carrying the SNP has been duplicated producing an insertion in some other place in the genome. The marker involved in this duplication is labeled M* and can be thought of as having been identified as a “candidate” marker given its non-Mendelian behavior in terms of heterozygote calls (see Materials and Methods) while M_L_ and M_R_ correspond to its flanking Mendelian markers that are thus *not* part of the duplication. The region where the M* locus was initially mapped will hereafter be referred to as the “reference locus”. In Fig. 1A we assume that only Parent 1 carries the duplication and this duplicate copy has the allele of that same parent. For the purposes of the figure, we only represent M* in this duplication but other markers can very well be implicated too and if this is so one has even more evidence that there is a CNV. After crossing these two homozygous parents to produce an F1 individual, meiosis of the F1 leads to gametes that may shuffle the alleles of the parental chromosomes. In the case of a DH population, these gametes are used to produce diploid plants whose genomic content is that of a gamete but simply doubled. For the situation depicted in Fig. 1A where we focus on the reference locus and the duplication, the (gametic or DH) associated segregation patterns fall into 4 categories. Assuming that these two loci are on different chromosomes (or far enough away from each other on the same chromosome), the genotyping of these plants will generate a call for M* that will be “A” 50% of the time, “B” 25% of the time and “H” 25% of the time. Thus the marker will be detected as anomalous (non-Mendelian) in this mapping population, having too many “H” calls, and this is the simplest situation for which our method allows one to map the second locus. As indicated in Fig. 1A, we introduce the associated 3-marker genotype classes based on the alleles arising for the M_L_, M*, and M_R_ markers. The CNV situation depicted in Fig. 1A will lead to a characteristic signature when considering the genome-wide allele frequency profiles. To illustrate this, we simulated a DH population with the same characteristics as GABI, taking a marker M* from chromosome 4 (specifically marker PZA-000492026) and then we duplicated it onto chromosome 5. The resulting allele frequency profiles are displayed in Fig. 1B. To construct the profiles, we first assigned the individuals of the simulated population to one of the 6 classes defined via the 3-marker genotypes ML M* MR. There are 6 classes because M* can be A, B or H while we impose ML and MR to be of the same parental type because in practice these markers are very close on the chromosome (because of that proximity, almost all individuals in the population will satisfy the imposed property and so in practice this restriction serves really to filter out cases that have been improperly mapped). Then for each class of individuals, we determine the allele frequency of all the Mendelian markers genome-wide (0 means only the B allele arises for the considered marker, 1 means only the A allele arises, *cf*. Materials and Methods), and plotting these leads to the allele frequency profiles as displayed in Fig. 1B. The x-axis is the cumulated genetic position for each of these Mendelian markers. Also displayed are the corresponding smoothed frequency curves as well as the allele frequency obtained without separating the individuals into the 3-marker genotype classes (dashed black curve). In this example the BHB curve has a peak (pointing down) on chromosome 4 as expected (the reference locus for M*) but also a second peak pointing up on chromosome 5. This peak is corroborated with that of the BBB curve (down) at that same position. We can thus say that the BHB and BBB curves together provide strong evidence for a 2:1 CNV of the A/B_A/-type, where the reference locus is normal (A/B; parent 1 having allele A and parent 2 having allele B) while the second locus involves a duplicate copy (A/-; carried by parent 1 only and where the copy has the allele of Parent 1 for the reference marker M*; see detailed explanation of the encoding in Materials and Methods). In such a notation, four different possible 1:2 or 2:1 CNV types are enumerated in the form A/B_-/A, A/B_A/-, A/B_-/B, and A/B_B/-.

**Figure 1.**
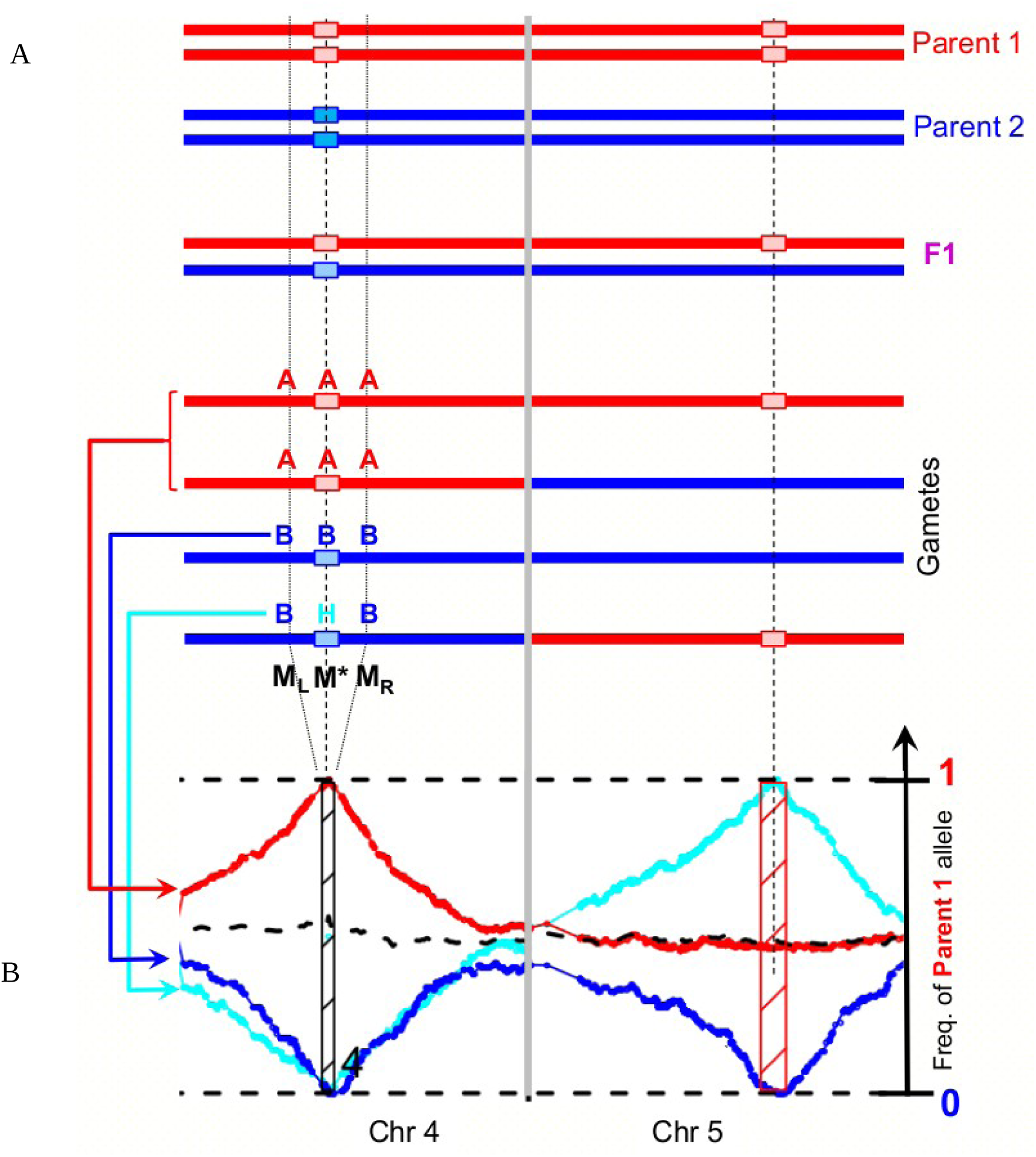
Consequences of parent-specific locus duplication on allele frequency profiles. A: Duplication in one parent leads to apparent heterozygotes in the gametes for the non-Mendelian marker M*. B: simulated genome-wide allele frequency profiles using subsets of individuals belonging to the three-markers genotype classes AAA, BBB, or BHB at markers ML, M*, and MR (ML and MR being the Mendelian markers flanking M*; see text). Such profiles reveal the loci involved in duplications. The allele of parent 1 is called “A”, the allele of parent 2 is called “B”, heterozygotes are called “H”, and missing data are called “-”. Each curve shows the frequency of the allele “A” along the genome (X-axis indicates cumulated genetic positions), when considering different subsets of individuals of the population as follows: cyan dots and curve for individuals (denoted “BHB”) genotyped “H” at the candidate marker M* and “B” on both non-candidate flanking markers ML and MR indicating the allelic context of the region, and similarly red for “AAA” individuals, dark blue for “BBB” individuals. Hatched rectangles indicate the estimated confidence intervals on the position of the detected loci involved in the event. The rectangle is black for the reference locus (see text) and red for the secondary locus. Dots represent values of individual markers and associated curves show the result of the smoothing procedure used to detect the peaks. Lastly, the black dashed line indicates the frequency of “A” allele based on all individuals of the population.

Analysis of the GABI data leads to many markers M* compatible with scenarios like that of Fig. 1 or their analogs under parental or allele exchange. For instance in Fig. 2A we show the profiles for a case that was detected as a 1:2 CNV with the duplicated locus within parent 2 but carrying the allele A. For completeness, we show further examples in Supplementary Figure S1 to cover all four types of 1:2 or 2:1 CNVs. In all these cases, the hashed rectangles at the peaks delimit the regions where the software localized each of the two loci by using the profile shapes in the neighborhood of these peaks (see Materials and Methods). Furthermore, to add credence to the different CNV claims when analyzing the data, we systematically provide simulations to determine the *expected* profiles under the hypothesis of the inferred scenario. Specifically, our software produces a simulated segregating population using the same number of individuals and the same marker positions as in the experimental data set but including one or more duplicate copies of M* at the position(s) predicted by the scenario (see Materials and Methods for a detailed explanation). Fig. 2B thus shows the expected profiles in the 1:2 CNV inferred from Fig. 2A while Supplementary Figure S1 includes the simulation for each of the four types of 1:2 or 2:1 CNV. If the result of a simulation shows patterns of peaks very close to the experimental ones, then one can have high confidence in the proposed CNV hypothesis.

**Figure 2.**
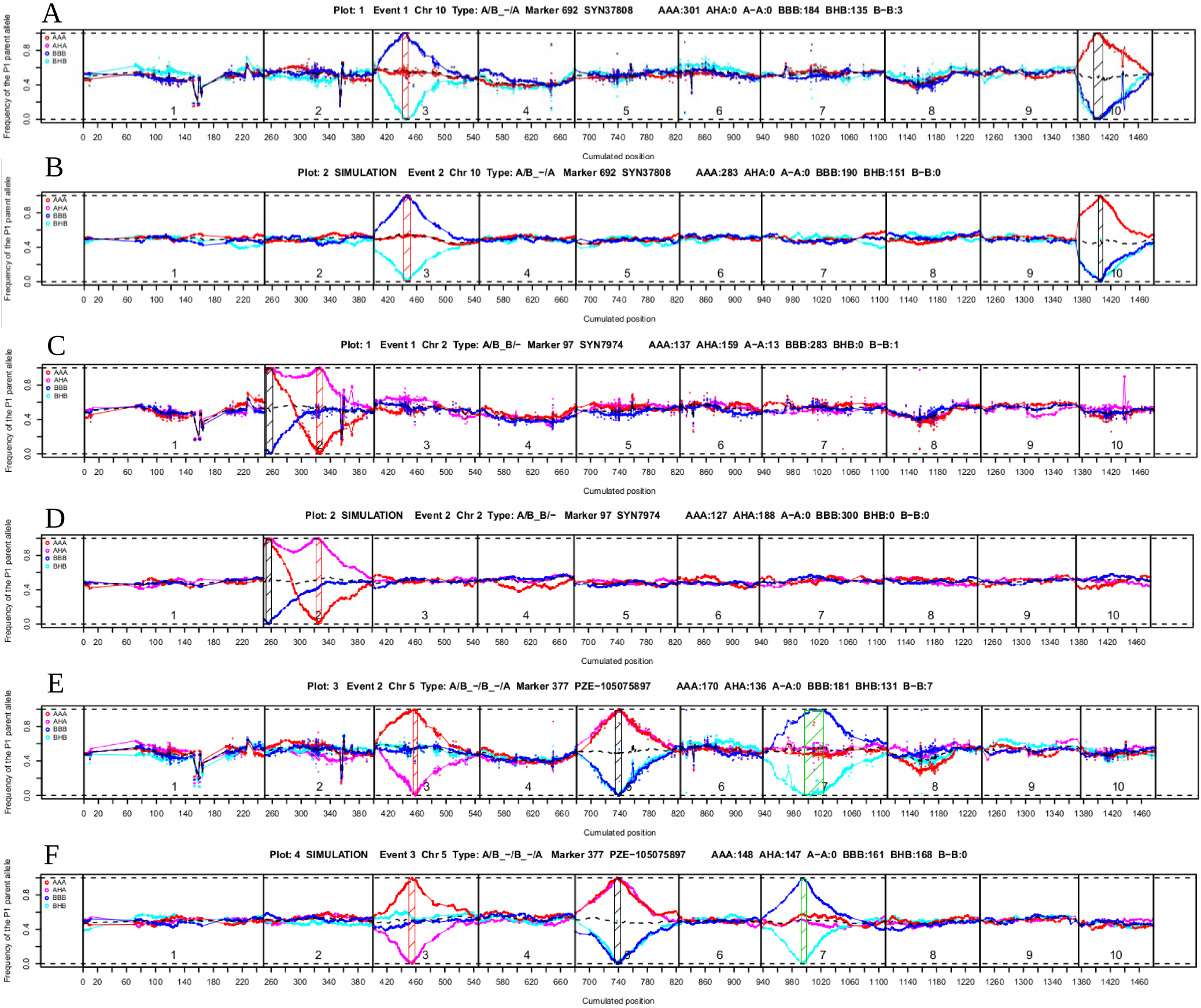
Examples of signatures of events involving two or three loci. Data are from the Maize doubled-haploid population GABI. Dots and curves have the same meaning as in Figure 1. Panels A, C, and E show experimental profiles for respectively a 1:2 CNV event with both copies on different chromosomes, a 2:1 CNV event with both copies on the same chromosome, and a 3-locus event with copies on three different chromosomes. Panels B, D, and F show simulation results reproducing the CNV situation inferred from panels A, C, and E respectively (see text). The allele of parent 1 is called “A”, the allele of parent 2 is called “B”, heterozygotes are called “H”, and missing data are called “-”. Each curve shows the frequency of the allele “A” along the genome (X-axis indicates cumulated genetic positions), when considering different subsets of individuals of the population as follows: pink dots and curve for individuals (denoted “AHA”) genotyped “H” at the candidate marker and “A” on both non-candidate flanking markers indicating the allelic context of the region, and similarly cyan for “BHB” individuals, red for “AAA” individuals, dark blue for “BBB” individuals. Curves generated by the software for classes based on missing data (light grey for “A-A” individuals, and black for “B-B” individuals) were hidden here for better clarity of the profiles. Hatched rectangles indicate the estimated confidence intervals on the position of the detected loci involved in the event. They are black for the reference locus (see text) and red for the secondary locus (or red or green for the two secondary loci in the case of the 3-locus event in panels E and F). The name of the candidate (non-Mendelian) marker considered is given in the header of each panel, as well as numbers of individuals counted for each three-locus genotype class.

The computer generation of all the profiles presented in this paper were obtained using the R package CNVmap available as Supplementary File S1.

#### The two loci of 1:2 or 2:1 CNVs sometimes arise on the same chromosome

The examples just shown had the duplicated locus on a different chromosome from the reference locus, but we also found other cases where the two peaks lay on the *same* chromosome. For illustration, we show such a case in the GABI population in Fig. 2C, the candidate marker being SYN7974. As in the previous case, our software produces a plot of the allele frequency profiles both for the experimental data and for a simulated data set given the inferred scenario which for Fig. 2C is A/B_B/-, both loci lying within chromosome 2. Clearly, the simulated profiles that are shown in Fig. 2D have all the qualitative properties seen in the experimental data, providing strong evidence that the parent 1 really does have a duplication of the region containing the SYN7974 marker and that the corresponding allele is that of parent 2. Compared to the case where the two loci are on different chromosomes, the expected proportion of individuals carrying the heterozygote signal is reduced: instead of the theoretical 25%, it is *r*/2 where *r* is the recombination rate between the two loci. Whenever these two loci are very close, the number of such recombinant individuals will be low and so it will be much more difficult to argue that there is a real CNV *vs* simply a few genotyping errors.

#### Allele frequency profiles reveal different types of 1:3 or 3:1 CNVs

Our software is set up to detect any number of peaks in the allele frequency profiles. Thanks to this feature, we found multiple cases where there were three separate loci. We illustrate such a situation in Fig. 2E in the GABI population for marker PZE-105075897, where the reference locus is on chromosome 5, and two additional copies were detected on chromosomes 3 and 7. The software automatically identifies this as an A/B_-/B_-/A, which means that the parent 1 has no additional copy while the parent 2 has two additional copies: one on chromosome 3 with allele “B” and one on chromosome 7 with allele “A”. Note that the peaks localizing these additional copies arise from the 3-marker genotypic classes AAA and AHA on chromosome 3, and the 3-marker genotypic classes BBB and BHB on chromosome 7. Again, to have a high level of confidence that the patterns observed have been properly interpreted, one can compare with the results of a simulation as was done in the previous figures. The result of simulating the triplication inferred from Fig. 2E is displayed in Fig. 2F, showing that the 1:3 CNV hypothesis is indeed strongly supported by the experimental data because of the high similarity between the Figs. 2E and 2F. Note that it is possible to show that the theoretical frequencies of the AAA, AHA, BBB, and BHB genotypes are 1/4, and of course this result agrees with what we see at the top of Fig. 2F and is not far from what is observed in the experimental case. In Supplementary Figure S2 we display similar cases but arising this time in the WHEAT population, corresponding to three-locus events of the types A/B_-/A_-/A, A/B_A/-_-/A, A/B_A/-_A/-, A/B_B/-_-/B, and A/B_B/-_B/-. Note that in all these last cases for which one of the alleles arises solely at the reference locus, the additional loci are identified only through one of the 3-marker genotypic classes, the classes having heterozygotes giving rise to enhanced frequencies at those two loci but not reaching the 100% value (see Supplementary Figure S2). The reasons one has enhancement but not a saturated peak or trough is that the constraint of capturing the multiple-copy allele can be satisfied at *either* of the two additional loci.

#### Missing data also can provide convincing signatures for 1:2 or 2:1 CNVs in the presence of systematic genotyping errors

In Fig. 3A we show a case arising within the GABI population, constructed based on the M* marker PZE-104096422. The patterns of the profiles resemble those of a 2:1 CNV except that the “BHB” profile is replaced by a similar one labeled “B-B” where the “-” means the call of the M* marker is “missing data”. We denote this case A/B_A/-|Hm to indicate that “H” calls were erroneously and systematically turned into missing data. Because this situation happens many times in the GABI population, we investigated a few cases in detail by examining the fluorescence data, the calls and the cluster file used with the Illumina array data. Given the two clouds of points produced from the fluorescence data for the cases of A and B calls, we find that the “-” calls typically correspond to a region that lies between those two clouds. Thus it is plausible that these cases, called as “-”, are in fact H, the discrepancy being due to a miscalibration of the cluster file. Based on this observation, we implemented in our code a procedure whereby the peaks of missing data detected in the allele frequency profiles could be interpreted as being due to such a “rule” according to which some proportion or even all of the H calls of M* become transformed into “-” calls (see details in Supplementary File S5). If only a fraction becomes transformed, both the BHB and B-B profiles provide a peak but if all H calls are transformed into “-” calls as seems to be the case in Fig. 3A, then the BHB curve will be absent. This reconsideration of the data in effect introduces a way to overcome the technical problem of inadequate cluster files that we observed to arise in the GABI population data. We also implemented the possibility of applying that transformation rule on *simulated* data, dependent on the probability of transforming an H call into a “-” call. That probability was estimated from the data. The resulting simulated profiles based on the inference of a 2:1 CNV in Fig. 3A are displayed in Fig. 3B, showing an excellent agreement between theory and experiment. Furthermore, this new class of events leads us to define a signature to be “strong” if each locus that is inferred to be involved in a CNV is identified by at least one peak from a 3-marker genotypic class without missing data, i.e., AAA, AHA, BBB or BHB. As seen in Figs. 3A and 3B, the locus carrying the putative duplication is identified by a peak for B-B but also by a peak for BBB and thus this event is associated with a strong signature. Clearly, all of the events illustrated in the previous sections correspond to strong signatures. We now move on to a more complex case where the second locus contains peaks but only for missing data and thus corresponds to a *weak* signature. As motivation for this more complex case, note that a miscalibration of the cluster file may be sufficiently severe that the H genotypes are called not only as “-” but also as either A or B. If that is the case, the peak in the previous case arising for the BBB 3-marker genotypic class no longer reaches the allele frequency zero at the second locus because some of the individuals contributing to the BBB class correspond in fact to BHBs. The result of these miscalls is the increase of the BBB frequency up from zero and thus the more or less disappearance of the BBB peak. Although the second locus of the CNV can be localized by the B-B curve, it is no longer detected *via* the BBB curve and this can raise some doubts as to the veracity of a 1:2 or 2:1 CNV interpretation. In Fig. 3C we show such a case, produced from the GABI data set for the candidate marker PZE-104127025. Because we have implemented the rule of transforming H calls into both “-” and “B” calls, the software detects this event and classifies it as a 2:1 CNV, denoted as A/B_A/-|HmHb to reflect the fact that “H” calls were erroneously systematically turned into either missing data or “B” calls. For these types of events also, our software provides a simulation of what should be expected under the corresponding hypothesis, estimating from the data the error rates turning H calls into “-” or into B; Fig. 3D shows the corresponding result, from which one may conclude that probably the weak signal in Fig. 3C is indeed indicative of a A/B_A/-|HmHb event.

**Figure 3.**
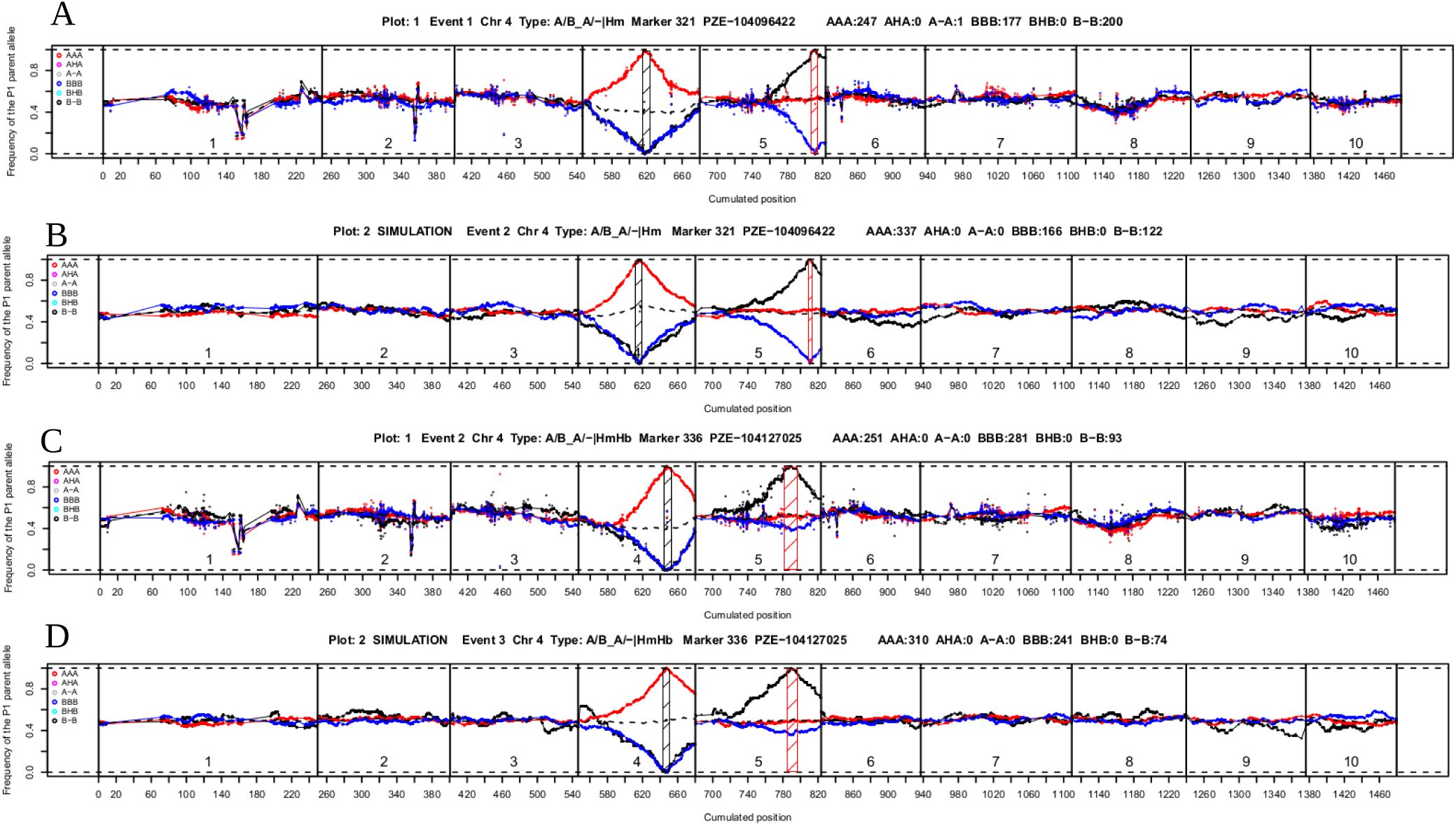
Examples of profiles showing characteristic signatures of CNVs in the presence of systematic genotyping errors. Data are from the GABI population. Dots and curves have the same meaning as in Figure 1. Panel A shows a typical “strong” signature, with a 2:1 CNV event in the case where “H” calls of the candidate marker were systematically called missing data (“-”). Panel C shows a typical “weak” signature, where a 2:1 CNV event in the case where “H” calls of the candidate marker were systematically called either as missing data (“-”) or as “B” in non-zero proportions. The software provides estimated systematic error rates for each such candidate. Panels B and D show simulation results reproducing the CNV situation inferred from A and C respectively (see text). The allele of parent 1 is called “A”, the allele of parent 2 is called “B”, and missing data are called “-”. Each curve shows the frequency of the allele “A” along the genome (X-axis indicates cumulated genetic positions), when considering different subsets of individuals of the population as follows: pink dots and curve for individuals (denoted “AHA”) genotyped “H” at the candidate marker and “A” on both non-candidate flanking markers indicating the allelic context of the region, and similarly cyan for “BHB” individuals, red for “AAA” individuals, dark blue for “BBB” individuals, grey for “A-A” individuals, and black for “B-B” individuals. Hatched rectangles indicate the estimated confidence intervals on the position of the detected loci involved in the event. The rectangle is black for the reference locus (see text) and red for the secondary locus. The name of the candidate (non-Mendelian) marker considered is given in the header of each panel, as well as numbers of individuals counted for each three-locus genotype class.

### Analyses of all events across all 3 mapping populations

We now move on and summarize what comes out of the analyses of each population when considering all of the corresponding candidate markers. Some characteristics of these populations, in particular their size and number of markers, are given in TABLE 1.

**TABLE 1.**
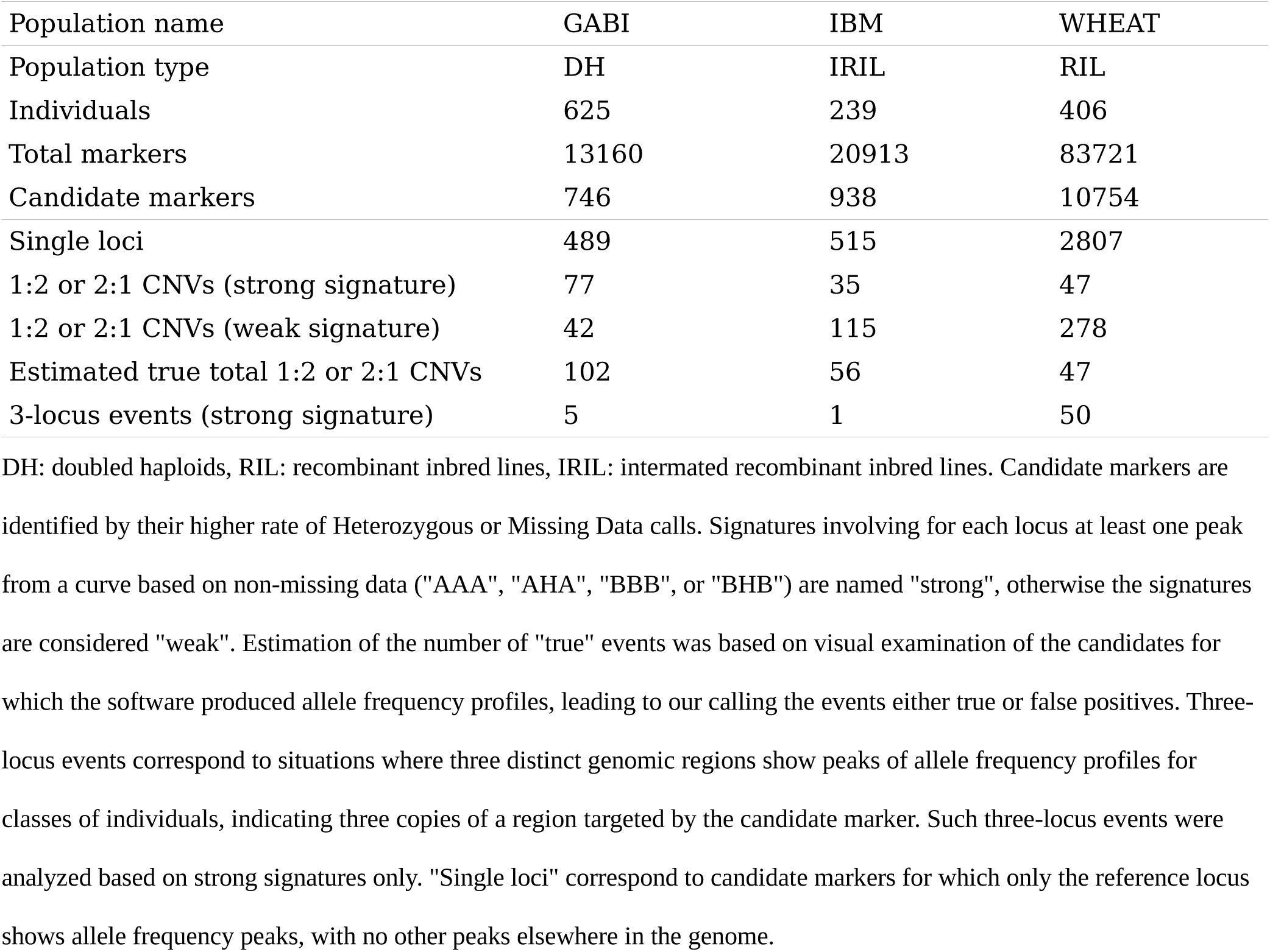
Results of automatic CNV detection in three different mapping populations.

| Population name | GABI | IBM | WHEAT |
| --- | --- | --- | --- |
| Population type | DH | IRIL | RIL |
| Individuals | 625 | 239 | 406 |
| Total markers | 13160 | 20913 | 83721 |
| Candidate markers | 746 | 938 | 10754 |
| Single loci | 489 | 515 | 2807 |
| 1:2 or 2:1 CNVs (strong signature) | 77 | 35 | 47 |
| 1:2 or 2:1 CNVs (weak signature) | 42 | 115 | 278 |
| Estimated true total 1:2 or 2:1 CNVs | 102 | 56 | 47 |
| 3-locus events (strong signature) | 5 | 1 | 50 |

DH: doubled haploids, RIL: recombinant inbred lines, IRIL: intermated recombinant inbred lines. Candidate markers are identified by their higher rate of Heterozygous or Missing Data calls. Signatures involving for each locus at least one peak from a curve based on non-missing data ("AAA", "AHA", "BBB", or "BHB") are named "strong", otherwise the signatures are considered "weak". Estimation of the number of "true" events was based on visual examination of the candidates for which the software produced allele frequency profiles, leading to our calling the events either true or false positives. Three-locus events correspond to situations where three distinct genomic regions show peaks of allele frequency profiles for classes of individuals, indicating three copies of a region targeted by the candidate marker. Such three-locus events were analyzed based on strong signatures only. "Single loci" correspond to candidate markers for which only the reference locus shows allele frequency peaks, with no other peaks elsewhere in the genome.

#### The GABI DH population

This population is very large (625 individuals, *cf*. TABLE 1) and there is no issue of residual heterozygosity coming from incomplete fixation because of the genome doubling. Given the thresholds used to classify the markers into the classes Candidate, Mendelian and “Other” (*cf*. Supplementary Table S1), we obtain 746 candidate markers (out of the total 13160). The software automatically analyzes the profiles associated with these markers to identify peaks and corresponding loci. Of these markers, 489 (a wide majority) lead to profiles involving a single locus (TABLE 1). In effect, these markers were assigned to the class Candidate because of technical problems with the genotyping, producing too many H or “-” calls, presumably because of some issues with the cluster file calibration rather than a presence of CNVs. One such case arises for marker SYN12874; it is presented in Supplementary Figure S3A and detected as “single locus” by our software. However, the remaining candidate markers lead to profiles having at least two loci (see TABLE 1).

About half of such multilocus cases are identified by the software as being proper 1:2 or 2:1 CNVs but their signatures are split between strong and weak. Visual examination of these profiles allowed us to validate or not these events, leading to an estimate of a total of 102 true 1:2 or 2:1 CNVs in this data set (*cf*. TABLE 1); 17 events were not validated by this visual inspection (putative false positives), corresponding to 16 with weak signatures and only one with a strong signature (Supplementary Table S2). Furthermore, in TABLE 2 we give the number of 1:2 or 2:1 CNVs found for each of the 4 possible cases, A/B_-/A, A/B_A/-, A/B_-/B and A/B_B/-. In a duplication-divergence scenario, one could hypothesize that a distant ancestor of one of the individuals formed an additional copy that subsequently diverged by mutation at a single base (Ohno 1970; Lynch and Conery 2000). In such a scenario, one might naively expect enrichments of A/B_A/- and A/B_-/B over the other two classes.

**TABLE 2.** Number of each type of event found in three segregating populations.

| Population name | GABI | IBM | WHEAT |
| --- | --- | --- | --- |
| A/B_-/A | 21 | 0 | 3 |
| A/B_A/- | 17 | 13 | 12 |
| A/B_-/B | 18 | 22 | 15 |
| A/B_B/- | 20 | 0 | 17 |
| A/B_-/A_-/A | 0 | 0 | 5 |
| A/B_A/-_A/- | 0 | 0 | 1 |
| A/B_-/B_-/B | 3 | 1 | 0 |
| A/B_B/-_B/- | 0 | 0 | 9 |
| A/B_-/A_A/- | 0 | 0 | 0 |
| A/B_A/-_-/A | 1 | 0 | 16 |
| A/B_-/A_-/B | 0 | 0 | 0 |
| A/B_-/B_-/A | 1 | 0 | 0 |
| A/B_-/A_B/- | 0 | 0 | 0 |
| A/B_B/-_-/A | 0 | 0 | 0 |
| A/B_A/-_-/B | 0 | 0 | 0 |
| A/B_-/B_A/- | 0 | 0 | 0 |
| A/B_A/-_B/- | 0 | 0 | 0 |
| A/B_B/-_A/- | 0 | 0 | 0 |
| A/B_-/B_B/- | 0 | 0 | 0 |
| A/B_B/-_-/B | 0 | 0 | 19 |

Number of events found for each category of 1:2 or 2:1 CNVs (upper part) or 3-locus events (lower part) in two Maize populations (GABI and IBM), and one wheat population (WHEAT). Every event was automatically detected by the software, and also visually checked by looking at the corresponding allele frequency profiles. Each category is encoded as a string of 2 to 3 groups of 3 characters each, separated by an underscore. The first group is always encoded "A/B" and indicates the reference locus, located at the position where the candidate marker was initially mapped. Further groups indicate other copies of the region targeted by the candidate marker. For all groups (loci), the letters just before and just after the slash represent respectively the haplotypes of the first parent (alleles denoted "A"), and of the second parent (alleles denoted "B").

However, in view of the numbers in TABLE 2, there is no such enrichment as all four classes have occurrence numbers of similar magnitude, a point that will be justified in the discussion. It is possible to further analyze the 1:2 or 2:1 CNVs by considering the separation into strong and weak signatures. Supplementary Table S1 gives the numbers for all types of automatically detected events. As indicated in Supplementary Table S2, many events with weak signatures are *not* validated by our visual inspection, so it is best to concentrate on the events providing strong signatures.

Of the other multilocus events, only 5 have a strong signature involving three or more loci (*cf*. TABLE 1). These 5 events belong to just a few of the different types with respect to allelic content as shown in TABLE 2; although it may be tempting to argue for an enrichment of the A/B_-/B_-/B type, the numbers are very small so it is not useful to go into such speculations.

Lastly, the software identifies 131 events in which there are multiple loci but with patterns of profiles that are “unknown” because they differ from those produced by CNVs in the list provided in Supplementary Table S1. Might some of these cases reveal true CNVs that are novel compared to what we considered so far? To get some insight into that possibility, we examined the corresponding profiles. Some cases provide no compelling evidence for a CNV, the profiles are simply very noisy and peaks may be presumed to be non-significant. For other cases, as in Supplementary Figures S3B and S3C, there is clearly an additional locus but the profiles are not as expected from our list of standard 1:2 or 2:1 CNVs. We also find cases of more than one additional locus as in Supplementary Figures S3D and S3E, where again the signature is not compatible with any of our standard 3 locus events or extensions.

#### The IBM IRIL population

Similar statistics for the IBM RIL population are given in TABLE 1 and TABLE 2, and in Supplementary Tables S1 and S2 and differ mainly quantitatively when compared to the GABI population. Nevertheless, from the conceptual point of view, the main new feature when going from GABI to IBM is the presence of residual heterozygosity in Mendelian markers. Indeed, since IBM is an intermated RIL population, fixation can be incomplete because either not enough generations of selfing have been applied or because there is a selective force impeding the homozygous state, situations that do not occur with doubled haploids. But once appropriate thresholds are set for the minimum number of heterozygote calls to select a marker as a candidate, our method was also efficient to discover CNVs in that population. As with the GABI population, the rate of true over false positives was extremely high (100% here, see Supplementary Table S2) when considering events with strong signatures. On the other hand, events with weak signatures gave a higher proportion of false positives in IBM as compared to GABI. Finally, there was only a single event associated with three loci. It should be noted that both IBM and GABI populations were genotyped with the same Illumina Maize SNP50 array, and deal with the same species, which is consistent with the qualitatively similar results obtained.

#### The WHEAT RIL population and importance of homeologues

The case of the WHEAT population is *a priori* quite different from the two first populations because bread wheat is hexaploid and also because the population was genotyped with a different SNP array. Not so surprisingly, we observed quite different results (see TABLE 1 and TABLE 2, and Supplementary Tables S1 and S2). This population is quite large (406 individuals, *cf*. TABLE 1) and it has far more markers (83721) than the other populations, justifying that the number of candidate markers, 10754, is also much higher (see Supplementary Table S1 for the thresholds used to define Candidate and Mendelian markers). In contrast to the other populations, the great majority of these candidates do *not* give rise to any profile, even the single-locus one, which is indicative of potential mapping problems. Nevertheless, a fair fraction of the candidates does give profiles. A large number of these, 2807 specifically, are identified as having just the reference locus and thus are not of interest. Such quite frequent cases are expected in WHEAT as in IBM because residual heterozygosity produces false candidates.

Of the remaining candidates, some are identified by the software as associated with 2 or more loci. For the events detected as being of the 1:2 or 2:1 CNV type, 47 have a strong signature and 278 have a weak signature. Validation by inspecting all of these events suggests that only the strong signatures provide true positives. In TABLE 2 we give the number of 1:2 or 2:1 CNVs found for each of the 4 types, A/B_-/A, A/B_A/-, A/B_-/B, and A/B_B/-. Though these classes are less balanced than in the GABI population, the evidence for enrichment of particular classes is not very strong. Supplementary Table S1 gives the numbers for all types (including thus the rules to take into account genotyping errors) of automatically detected events. However, as indicated in Supplementary Table S2 and previously mentioned, the events with weak signatures are not validated by our visual inspection, so it is best to focus on the events with strong signatures only.

This brings us to the strong signatures for events involving three or more loci (*cf*. TABLE 1). The types of these events are given in TABLE 2. Clearly the main types seen have one allele in three copies and the other in a single copy. It is relevant here to recall that wheat is a hexaploid which contains three genomes (A, B, and D) with seven chromosomes each. We found 20 cases where the three copies are located on three homeologous chromosomes (e.g. in Supplementary Figures S2C and S2E on chromosomes 2 and 6), 8 cases with two copies on two homeologous chromosomes and the third one on a different (non-homeologous) chromosome (e.g., in Supplementary Figures S2A and S2I), and finally 22 cases where the three loci are on three non-homeologous chromosomes (e.g., in Supplementary Figure S2G). Although enrichment amongst homeologues is expected, it is appealing to have it come out from our automatized software.

#### Sequence-based validation of candidates with strong but not weak signatures for IBM

CNVmap provides candidate CNVs and predictions for the associated loci. In the case of the IBM population, the parental genomes are fully assembled and so our predictions can be checked by searching in those reference genomes for multiple occurrences of the specific sequences flanking the candidate SNPs *via* BLAST (see Methods for details). The results of those analyses are as follows. First, concerning events having strong signatures, the majority of the predictions are validated. Specifically, of the 13 predicted 2:1 CNVs (A/B_A/- or A/B_B/-; two copies in parent B73), all are validated, while of the 22 predicted 1:2 CNVs (A/B_-/A or A/B_-/B; two copies in parent Mo17), 12 are validated (Fig. 4). Not surprisingly, when testing the hypothesis H_0_ of no CNV in these strong signature events, all but one of the *p*-values (for events validated or non-validated by BLAST) were below 0.05 and most of them were below 10^-3^ (Fig. 4, see Materials and Methods for the calculation of these *p*-values). Second, concerning events having weak signatures, essentially none of them are validated; furthermore, Fig. 4 shows a broad distribution of *p*-values, calling in doubt the credibility of these weak candidate CNVs.

**Figure 4.**
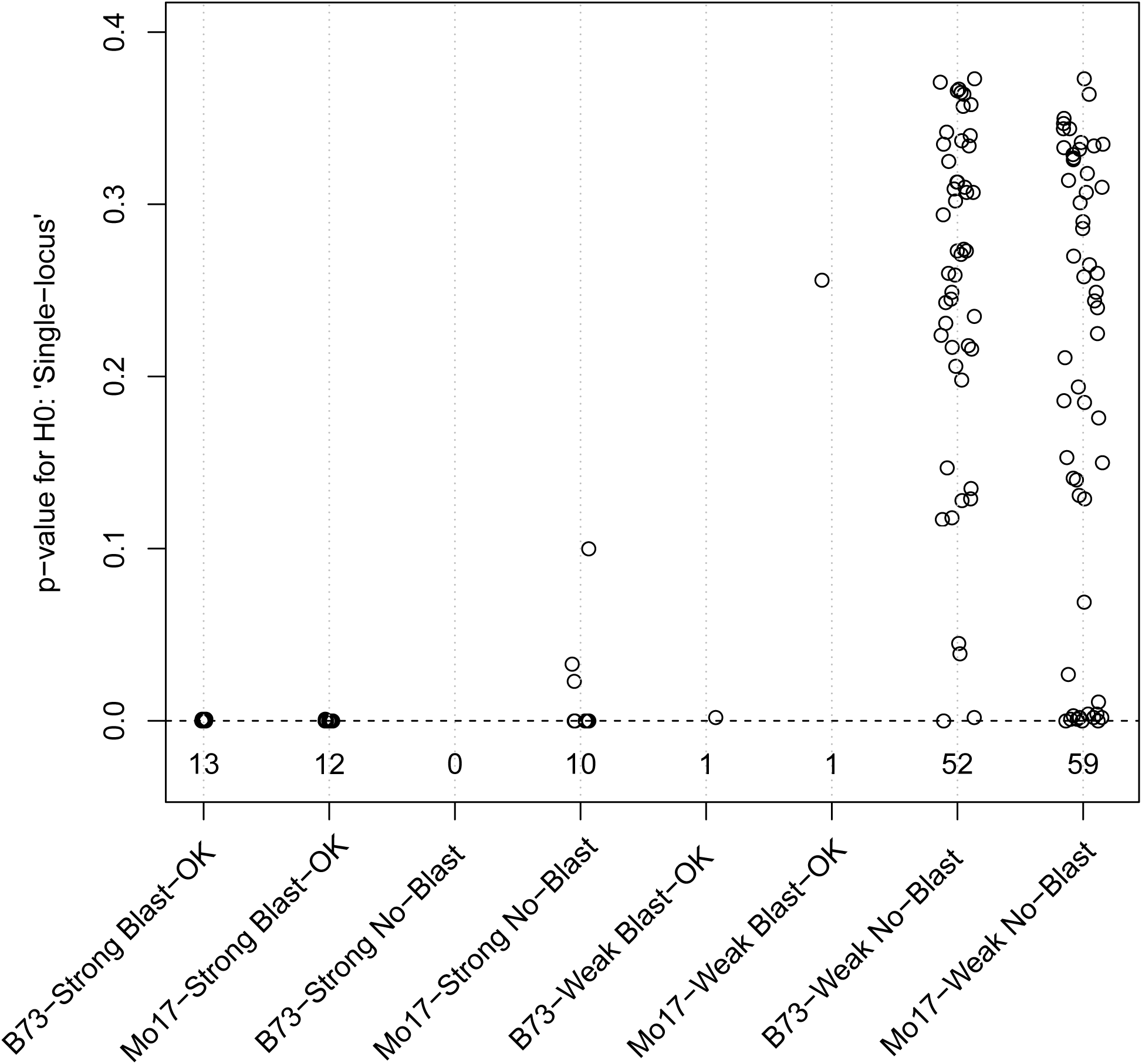
Validation of CNVs found in the IBM population. All 1:2 and 2:1 CNVs found in the IBM population (obtained from the cross B73xMo17), based on Strong or Weak signatures, were submitted to two different types of validation (see Materials and Methods): (1) a *p*-value (Y-axis) was computed using 1000 simulations for the H_0_ hypothesis: “the marker M* is present as a single copy in both B73 and Mo17 parents”, and (2) the presence of the second copy in the reference genome of the expected parent (B73 for 2:1 CNVs and Mo17 for 1:2 CNVs) was checked using BLAST search against whole genome sequence assemblies of both parents. On X-axis, events are denoted ‘Blast-OK’ or ‘No-Blast’ according to the success of the sequence-based validation, and ‘B73 (or Mo17)-Strong (or Weak)’ according to the types of events considered (strength of the signature and B73 or Mo17 having two copies). Numbers below the line Y=0 indicate the number of events in each category.

## Discussion

### An original method based on linkage to detect and genetically map CNVs

We presented a new method for revealing and genetically mapping copy number variations in bi-parental segregating populations made of homozygous individuals. The heart of the method is the fact that a marker participating to a CNV will lead to an excess of heterozygotes in the segregating population with associated signatures in genome-wide allele frequency profiles. We validated this method with maize doubled-haploid lines, maize intermated recombinant inbred lines, and wheat recombinant inbred lines, revealing CNVs even in these last two types of populations in spite of the presence of their residual heterozygosity. The approach does not involve any *a priori* knowledge about the type or location of the events, rather it is based on signatures in genome-wide allele frequency profiles, assuming that the individuals therein have been genotyped. Such genotyping might have been done for instance for detecting QTLs or for breeding purposes (as in genomic selection), and thus our approach can “piggy-back” *for free* after the production of such genetic material. In this context, our detection of CNV loci has a spatial resolution that depends on the local recombination rate, so the larger the population and the larger the recombination rate the better. Nevertheless, detecting the *existence* of duplicated loci and finding their approximate localization is relatively easy: 239 individuals are sufficient for the RILs (F6) we studied, and lower numbers can also give good results. The major limitation of our approach is that the duplicated loci cannot be too close to the original locus, and thus we cannot easily detect tandem duplications. Another requirement of course is that the markers be robustly ordered, so the quality of the genetic map is important. In usual genotyping arrays, SNPs have been included only if they were found to be exploitable on a reference panel, and thus SNPs with heterozygous signals have little chance of having been kept for inclusion on that array. As a result, we certainly strongly underestimate the real number of CNVs. Consequently, our approach, when used on data produced with SNP arrays, should not be considered as a way of surveying the *number* of structural variation events between two parents, but rather as a cost-free means of getting, for a subset of such events, detailed information on their nature and in particular the genomic locations of the associated copies.

### Different rate of success of sequence-based validations in B73 and Mo17 parents

In the IBM population, all 2:1 CNV events (with two copies in parent B73) were confirmed by BLAST analysis, whereas only 55% of 1:2 CNVs (with two copies in parent Mo17) led to successful validation. This difference may be due to a less good quality of the genome assembly of Mo17. Indeed, the quality of B73 assembly is most probably higher because that inbred line was chosen for the first Maize reference genome sequence, so more sequencing and assembly effort was dedicated to this line. Another explanation may be that the level of sequence divergence between B73 and Mo17 leads some loci to escape our BLAST search on Mo17 because the oligonucleotides used in the MaizeSNP50 Illumina array were designed based on the B73 reference sequence, but nevertheless the markers would still be able to hybridize on Mo17 DNA.

### Applicability to non-fixed populations

Our method is primarily based on the detection of apparent heterozygosity, so the presence of heterozygous loci due to incomplete fixation, as occurs in RIL populations, is expected to greatly lower the efficiency of detection. So we adapted the criterion for a marker to be a non-Mendelian candidate by enforcing its level of heterozygosity to be higher than a given threshold, thereby limiting the number of candidate markers to analyze. And in fact, the method proved to be sufficiently powerful to detect CNVs in recombinant inbred line populations corresponding to six generations of selfing.

### Detecting CNVs in the presence of systematic genotyping errors

We also found clear signatures of CNVs based on missing data. Typically, such missing data arise from technical systematic biases in the genotyping (e.g. systematic mis-calling of heterozygotes as missing data and/or as homozygotes), and thus can be put on a similar footing with the more standard signatures of CNVs such as A/B_B/-. Thus, in addition to being able to detect non-Mendelian SNPs in a genotyping array (in linkage analyses like QTL mapping, it is useful to remove them), our method is also able to reveal some flaws in the cluster files used for analyzing Illumina array data. Such cluster files, which determine the way fluorescence levels at two different wavelengths are converted into a genotype call, may be more or less appropriate depending on the genetic origin of the material being genotyped, or may be sensitive to some variations of the experimental conditions during the hybridization of the arrays. In our results, we could clearly demonstrate that a large number of markers suffered from such systematic genotyping errors transforming heterozygote calls into missing data and/ or into homozygotes.

### Most detected events correspond to four types of parent-specific duplications

For the two maize datasets studied, the vast majority of the events involve just two loci. Furthermore, most of these can be identified as parent-specific duplications, corresponding to the four types A/B_A/-, A/B_-/A, A/B_B/-, or A/B_-/B. Among them, A/B_A/- and A/B_-/B involve haplotypes with two copies of the same allele in the parent carrying the duplication. On the other hand, A/B_-/A and A/ B_B/- involve two different alleles at the additional locus in the haplotype of the parent carrying the duplication, so that parent -- although it is an inbred line supposed to be almost fully homozygous -- is expected to be genotyped as heterozygous for marker M*. We thus analyzed genotyping data obtained from the parents, and indeed, in all such cases, we observed heterozygous calls in the genotype data associated with the corresponding parent of the GABI population. From an evolutionary point of view, the latter situations (A/B_-/A and A/B_B/-) might seem to suggest a temporal order, namely a duplication followed by a divergence at the reference locus. However, it is just as likely that the divergence happened first and the duplication later: since the two loci are not tightly linked, recombination between them can very well produce the A/B_B/- haplotype starting with the A/B_-/B haplotype. Thus there is no *a priori* expectation that two of the four types of 1:2 or 2:1 CNV should be much rarer than the other two types, and this is in line with what comes out of the summary statistics as can be seen in TABLE 2 for GABI and WHEAT populations. However, in IBM, all 1:2 or 2:1 CNVs detected had the same allele on both copies for a given haplotype, which suggests that SNPs in the IBM mapping data set may have been selected to remove markers with heterozygous calls on the parents.

### The special case of wheat homeologous chromosomes

In the case of wheat which is a hexaploid species containing three diploid genomes, one has the further issue of homeologous chromosomes. Because these chromosomes have diverged from a common ancestor, the gene content is quite well conserved and chromosomes display good collinearity with limited rearrangements (Consortium (IWGSC) *et al*. 2018). A consequence of this is that SNPs may not necessarily be genome-specific and may therefore hybridize on two or three homeologous loci (Rimbert *et al*. 2018). Such similar sequences may generate signals of CNVs in the allelic profiles and so we asked the question of whether the duplicated loci we found in wheat were more often than expected on the homeolog. The analysis of the two- and three-locus events in our WHEAT dataset in fact shows a huge enrichment for favoring the homeologous chromosomes. Our method can thus provide a useful way to assess the level of genome-specificity of the SNPs of a given genotyping array, and help validating the selection of subsets of purely Mendelian markers.

### Only a tiny minority of the allelic profiles involve three or more loci in the maize populations

Because of the hexaploid nature of wheat, this plant was expected to reveal many triplication events if markers were not perfectly genome-specific, and this is actually what we found. On the other hand, in maize the ancient allotetraploid origin of the species is old enough for most markers to behave as single-copy, so one may expect far fewer three-locus events. And indeed, as can be seen from TABLE 2, there are some candidate markers that generate profiles with three loci but they are quite rare and arise mostly within the GABI population. This difference may be due to the GABI population being much larger, allowing our method to be more powerful on that data.

### Possible evolutionary scenarios for triplications

Some entangled events such as A/B_B/-_-/B may seem unexpected because they involve the allele from the opposite parent. However, just as we explained for the two-locus case, recombination can scramble the assignment of alleles and so a posteriori such events are not surprizing. But there is another possibility for justifying such an entangled CNV without appealing to recombination. Indeed, imagine that an ancestral triplication arose so that the allele B was present at all three loci. Parent 1 and Parent 2 may be identical by descent for that triplication for all of their homologues. If so, today’s situation can very well be due to subsequent divergence only: the divergence at the reference locus would produce a SNP while the divergence at the other two loci would be more severe, for instance corresponding to a deletion or appearance of other SNPs in the flanking sequences of the two other loci, thereby preventing the hybridization of oligonucleotides. Clearly such a scenario can also be responsible for entangled 1:2 or 2:1 CNVs.

### Conclusion

We developed an original *linkage-based* method to detect CNVs and genetically map the associated previously unknown copies from genotype data of segregating populations. Our software based on this method makes it possible to perform fully automatic mining of segregation data to extract a list of high confidence CNVs, including the detailed type of event and the genomic location(s) of the initially unknown locus or loci. It is thus a costless and easy way to generate additional added value from genotyping efforts initially dedicated to genetic map construction or QTL analyses. Because of its ease of use, our tool for detecting CNVs could be applied to other kinds of populations. First, going from bi-parental to multi-parental RILs as used in MAGIC (Dell’Acqua *et al*. 2015) or NAM (McMullen *et al*. 2009) populations should be straightforward, our computer program can be used as such for all biallelic SNPs. Second, it seems possible that CNVs could be detected by our approach when using the kinds of panels exploited in GWAS when the individuals in the panel are homozygous (e.g. inbred lines); the method would then correspond to searching genome-wide for associations between allele frequency and the particular 3-loci genotype (e.g., AHA) detected at a reference locus (that is for the non-Mendelian marker of interest and its two flanking markers). Such an approach, using a diversified panel, might in fact allow one to identify duplicated loci with a high level of resolution.

## Declarations

### Ethics approval and consent to participate

’Not applicable’

### Consent for publication

’Not applicable’

### Competing interests

The authors declare that they have no competing interests

### Funding

The research leading to these results has received funding from the French Government managed by the Research National Agency (ANR) under the “Investment for the Future” programs BreedWheat and Amaizing (project ANR-10-BTBR-03), from FranceAgriMer, French Funds to support Plant Breeding (FSOV) and from INRA. This work has benefited from a French State grant (LabEx Saclay Plant Sciences-SPS, ANR-10-LABX-0040-SPS), managed by the French National Research Agency under an “Investments for the Future” program (ANR-11-IDEX-0003-02) which funded the salary of KJ.

### Authors’ contributions

MF and OM conceived the method, developed the computer programs, analyzed and interpreted the results, and wrote the final manuscript. MF, KJ and OM worked on the computer program, performed analyzes on the data sets, and produced the first draft. EP, CK, and SM provided data sets. All authors read, edited and approved the manuscript.

## Acknowledgements

The authors are grateful to two anonymous reviewers for their comments which helped us to improve both the software and the manuscript. We also thank E. Bauer for fruitful discussions about the method, C. Schön, T. Mary-Huard and S. Nicolas for comments and advice, and to the CNV4Sel group for feedback during the development of our computational tool.

## List of Supplementary Material

**Supplementary Table S1.** Detailed statistics of the three populations GABI, IBM, and WHEAT

**Supplementary Table S2.** Assessment of rates of true and false positives by visual inspection

**Supplementary Figure S1.** Examples of allele frequency profiles showing characteristic signatures of four different types of 1:2 or 2:1 CNVs.

**Supplementary Figure S2.** Examples of allele frequency profiles for each type of three-locus events found in wheat.

**Supplementary Figure S3.** Examples of allele frequency profiles showing different situations where no CNV was identified by the software.

**Supplementary File S1.** CNVmap software as an R package (CNVmap12.tar.gz archive to open in R with the function install.packages()).

**Supplementary File S2**. Compressed (.zip) archive of GABI data sets to run CNVmap.

**Supplementary File S3**. Compressed (.zip) archive of IBM data sets to run CNVmap.

**Supplementary File S4**. Compressed (.zip) archive of WHEAT data sets to run CNVmap.

**Supplementary File S5**. Supplementary Methods: specification of all CNV signatures recognized by the software CNVmap.

**Supplementary File S6**. Installing with Rstudio. Particular procedure for some versions of R/Rstudio.

## Notes

#### Summary of Updates

The main new points in this revision are: - computation of a p-value was added to the software for the H0 hypothesis that there is no SV (the marker is single-locus on both parents) - sequence-based validation was carried out for the CNV candidates found in the IBM population based on the whole-genome sequence assemblies of the parents B73 and Mo17 - more stress was put on the fact that the novelty of the approach lies in the fact that our method not only detects but also genetically maps the CNVs

